# Ultrastructure of stemness and differentiated state in *Hydra* epithelial cells

**DOI:** 10.64898/2026.07.20.739505

**Authors:** Anna Seybold, Willi Salvenmoser, Kristian Pfaller, Stefan Redl, Michael W. Hess, Bert Hobmayer

**Author notes:** Correspondence: Anna Seybold,; Bert Hobmayer.

## Abstract

Epithelial cells in *Hydra* perform an unusual combination of functions: they divide continuously like adult stem cells while simultaneously executing the complex physiological tasks of differentiated epithelia. This challenges the traditional distinction between proliferative stem cells and terminally differentiated tissue, raising the question of how a single cell type integrates these opposing roles. Using electron microscopy, we examined morphological characteristics that define the stem-like and differentiated states of *Hydra*’s ectodermal and endodermal epithelial cells. Stemness is reflected by nuclear characteristics of active proliferation, including extensive euchromatin, large nucleoli, and the presence of nuage. However, differentiated epithelial cells exhibit strong apical-basal polarity, various endomembrane compartments for endocytosis and transport, specialized secretion mechanisms, and basal muscle processes with dense-core vesicles implicated in hormonal communication. Cryofixation improved ultrastructure preservation, elucidating the pleiomorphic configurations of complex intracellular channel systems traditionally presenting as singular vacuoles. This may shed new light on possible functions of this compartment. Taken together, *Hydra* epithelial cells combine ancient stem cell traits with highly specialized differentiated functions. This multifunctionality provides insight into the cellular organization of early-branching animals and suggests that multifunctional epithelia may represent an ancestral condition preceding the strict segregation of stem and differentiated cell lineages in bilaterians.

## Introduction

Epithelia represent an ancient and conserved tissue type, both, in multicellular animals (Metazoa) and in their eukaryotic, colony-forming relatives (Tyler 2003). Epithelia are defined by cells structurally and functionally polarized along an apical-basal axis and connected at their lateral membranes using cell-cell adhesion contacts, often located at an apical position along the lateral membrane. Metazoan epithelia produce an extracellular matrix at their basal face, and stably connect to this extracellular matrix using cell-matrix adhesion contacts. In bilaterians, epithelial cells are commonly regarded to represent cells in a differentiated, non-proliferative condition forming tissues with distinct physiological and structural functions. The capacity to renew and maintain these tissues during a bilaterian’s life span is thought to be restricted to small populations of epithelial stem cells with reduced functionality but potency for differentiation (Rinkevich et al. 2022). In contrast, in more ancestral, pre-bilaterian metazoans such as cnidarians and sponges, epithelial cells act in highly specialized functions in nutrition, physiology, water household, immunity, etc. and often simultaneously exhibit stem cell-like properties, challenging the exclusiveness of stem cell and differentiated states (Rinkevich et al. 2022).

*Hydra* is an ideal model for investigating the ancestral epithelial architecture, due to its simple anatomy and transparency, and to its experimental accessibility at the molecular level. It represents the classic cnidarian polyp body plan. Along its primary oral-aboral body axis, a differentiated head consisting of the mouth opening (hypostome) and a ring of tentacles to capture prey defines the oral terminal end. A differentiated foot to adhere to the substrate is located at the opposite aboral terminal end. In between, the gastric region exhibits a mostly undifferentiated part with large numbers of epithelial and interstitial stem cells, from which the head and foot cells derive by differentiation and movement. Traditionally, cnidarian epithelia are described as epidermal and gastrodermal layers emphasizing their physiological roles, separated by an extracellular matrix called mesoglea. In *Hydra*, the epithelial cells are termed ectodermal and endodermal myoepithelial cells, reflecting their function as musculature under neuronal control. In this study, we use the terms “ectodermal” and “endodermal” epithelial cells to follow the terminology commonly used in the *Hydra* literature. Ectodermal and endodermal epithelial cells constitute two independent adult stem cell lineages separated during late embryogenesis. A third and independent adult stem cell lineage, interstitial stem cells and their progeny, are interspersed in the two epithelial layers. This lineage produces neurons, nematocytes and gland cells, as well as gametes when polyps switch to sexual reproduction. Interstitial cells control polyp behavior, but play a minor role in morphogenesis and regeneration.

A central question in *Hydra* biology concerns the functional identity of its epithelial cells: decades of research suggest that *Hydra* epithelia cannot be easily placed into a specific category. In their function to shape the polyp body, epithelial cells interact with their freshwater environment (ectoderm), they act in prey capture (ectoderm and endoderm), digestion and nutrient uptake (endoderm), and mediate immunity and an associated microbiome. On the other side, epithelial cells display properties typically associated with embryonic tissues: continuous cell division, reduced G1 phase, active migration, dynamic body-axis remodeling, morphogenetic plasticity, and a renewing positional information system. By that, epithelial cells are the main drivers of shape generation and whole body regeneration. Together, these traits position *Hydra* as an “embryonic relic,” retaining cellular behaviors that may also resemble early metazoan epithelia. Several research articles highlight the plasticity and multi-functionality of *Hydra* epithelial cells at cellular and transcriptomic levels of analysis (Anton-Erxleben et al. 2009; Hemmrich et al. 2012; Buzgariu et al. 2015; Siebert et al. 2019; Cazet et al. 2023).

Here, we re-investigate the ultrastructural basis of stemness and differentiated states in *Hydra* epithelial cells. A large body of classic literature has described the major features of the two epithelial cell lineages (Hess 1961; Lentz 1966; Slautterback 1967; Haynes et al. 1968; Bibb and Campbell, 1973; Wood 1985; Campbell 1987; Rieger 1994). Now we show some of these features using modified techniques of fixation and tissue preparation, particularly the use of high-pressure freezing in order to allow the visualization of ultrastructure closer to its native state (Holstein et al. 2010; Böttger et al. 2012; Fraune et al. 2015)

By doing so, we are able to show that *Hydra* epithelial cells simultaneously exhibit elements of proliferating stem cells, including open chromatin, large nucleoli, and nuage at various cytoplasmic domains, as well as features of fully differentiated cells, such as specialized secretory and endocytic structures in apical and basal cell compartments, active transport systems including a large vacuolar channel system, various types of cell-cell and cell-matrix junctions, and basal muscle processes. By comparing these structural states, we add to the discussion how a single epithelial cell type integrates stemness with physiological specialization. This work provides insight into the evolution of epithelial plasticity and supports a view that multifunctional epithelia likely represent an ancestral condition in early animals.

## Materials and methods

### Animals and culture

*Hydra vulgaris* and *Hydra magnipapillata* polyps were used as standard wildtype strain. Transgenic Lifeact-GFP-expressing strains (Aufschnaiter et al. 2017) were used for imaging of macropinocytotic cups. Polyps were kept in asexual mass culture at a constant temperature of 18 °C as described (Hobmayer et al. 1997). They were fed daily with newly hatched Artemia nauplii and washed ∼8 h later. Experimental animals were starved for 24 h.

### Histology and light microscopy

Adult *Hydra* polyps were relaxed in 2% urethane in *Hydra* medium for 3 minutes and fixed in Carnoy’s fixative, a mixture of ethanol, chloroform and acetic acid, 6 + 3 + 1 respectively for 3 hours, dehydrated in ascending ethanol series and embedded in Technovit 7100 resin. Blocks were cut with an autocut 2040 (Reichert, Vienna). Three µm thick sections were stained according to (Richardson et al. 1960) and mounted. Images were taken with a Leica 5000DM light microscope (Wetzlar, Germany) using a Leica DFC 490 digital camera (Wetzlar, Germany).

### Preparation for transmission and scanning electron microscopy (TEM, SEM)

Chemical fixation: Animals were relaxed in 2% urethane in *Hydra* medium for 3 minutes and then fixed after Shigenaka et al. (1971) in 1/15 M PBS pH 7.2, 1 mM MgSO_4_, 0.1 M Sucrose, 3% glutaraldehyde and 1% OsO4 for 1 h on ice or after (Campbell 1987) in *Hydra* medium. After dehydration in a graded acetone series (50, 2×70, 90, 100, 15 min each), animals were incubated in a 1:1 mixture of acetone and EMBed 812 overnight, infiltrated for 3×1 h and polymerized in epoxy resin at 65 °C for 48 h.

Cryofixation: For TEM animals were high-pressure frozen, freeze-substituted with acetone (containing 0.5% OsO4 and 0.1% uranyl acetate) and embedded in Epon™ epoxy resin as described in (Holstein et al. 2010). For SEM high-pressure frozen animals were cracked with a pre-chilled scalpel or razor blade under liquid nitrogen, freeze-substituted, followed by critical point drying and sputter coating.

Semi-thin and ultra-thin sections (∼350 and 80 nm, respectively) were cut with a diamond knife (Diatome, Switzerland) on a Leica UCT ultra-microtome (Leica, Vienna), mounted on copper grids and optionally stained with lead citrate or hot ethanolic phosphotungstic acid (Locke and Krishnan 1971).

### Electron microscopy

TEM-samples were examined with a Zeiss Libra 120 energy filter transmission electron microscope (Zeiss, Germany) or a Philips CM120 transmission electron microscope (Philips, The Netherlands) equipped with a Tröndle 2x2 high speed camera (Tröndle, Germany) and ImageSP software (Tröndle, Germany), or a MORADA G1 digital camera (EMSIS, Münster, Germany) and iTEM-FIVE software (EMSIS), respectively. SEM-samples were examined with a ZEISS Gemini 982 field emission scanning electron microscope (Böttger et al., 2012). Image processing (including stitching) was carried out with ImageSP, iTEM-FIVE and Adobe Photoshop V.9.

## Results

### Epithelial cells express features of adult stem cells and functionally differentiated cells

*Hydra* epithelial cells display an intriguing multifunctionality: they maintain proliferative potential characteristic of adult stem cells, while at the same time performing the complex physiological roles of fully differentiated epithelial tissue. To understand how these opposing capacities coexist within a single cell type, we examined ultrastructural characteristics associated with stemness and differentiation.

Our observations reveal morphological features that define the dynamic state of epithelial cells and illustrate how structural specialization supports functional diversity. Figure 1 provides a light micrograph and a schematic overview based on electron microscopic data of the diverse morphological features, which will be characterized in ultrastructural detail later on in this study. *Hydra* epithelial cells display a clear set of traits distinguishing stemness from differentiation (Table 1). Stem-like cells are characterized by active cell-cycle stages (S-phase and mitosis) while remaining within the epithelial layer, large euchromatin-rich nuclei with prominent nucleoli, and the presence of nuage structures (Hobmayer et al. 2012; Juliano et al. 2014) associated with nuclear pores, mitochondria, or dispersed in the cytoplasm. In contrast, differentiated ecto- and endodermal cells exhibit defined epithelial polarity, including apical cuticle and junctional complexes, and show functional specialization such as apical granule secretion, channel/vacuole systems, phagocytic or nutritional uptake activities, basal muscle processes, and dense-core vesicle-communication. Layer-specific traits include cuticle formation and macropinocytosis in the ectoderm and cilia, microvilli, and nutrient uptake in the endoderm.

**Table 1:** Differentiation and stemness traits of epithelial cells: Criteria for stemness in both ecto- and endodermal cells are: s-phase and mitosis, nucleus, nucleolus and nuages. During s-phase and mitosis cells remain in the epithelial layer with intact SJ, but disassemble basal myoprocesses and contact to mesoglea (S-phase BrdU positive); The nuclear distribution of non-condensed to condensed chromatin is typical for stem cells. Nuages localize, as in stem cells, attached to nucleopores, mitochondria or “free swimming” in the cytoplasm. Criteria for differentiation in both ecto- and endodermal cells are: epithelial polarity, apical secretory granules, channel system/vacuoles, phagocytosis, basal muscle processes and dense core vesicle-communication. The cell polarity is defined by apical cuticle, or glycocalix and septate junctions (SJ), basal desmosome-like junctions (DLJ) between myoprocesses and hemi-DLJ with the mesoglea. Ectodermal differentiation traits are the five layered cuticle (glycosaminoglycans and several proteins) as well as the macropinocytosis. Endodermal differentiation traits are flagellae, microvilli with glycocalix as well as nutrition uptake.

| Criteria for | Stemness | Differentiation |
| --- | --- | --- |
| S-phase and mitosis (as a stem cell marker in *Hydra*) | ✓ |  |
| Nuclear morphology (large amount of non-condensed euchromatin) | ✓ |  |
| Nucleolus (large nucleolus as a marker for pluripotency, RNA metabolism) | ✓ |  |
| Nuage (as a clear marker for stemness) | ✓ |  |
| Epithelial polarity (epithelial configuration reflects differentiation state)<br>• Cuticle (ectodermal)<br>• Flagella, microvilli<br>• Junctions (apical SJ, basal DLJ & HDLJ)<br>• Basal muscle processes |  | ✓ |
| Transport<br>• Channel system/vacuoles<br>• Macropinocytosis (ectodermal)<br>• Nutrition uptake (endodermal)<br>• (Phagocytosis) |  | ✓ |
| Secretion<br>• Apical granules |  | ✓ |
| Hormone activity<br>• Dense core vesicle-communication |  | ✓ |

**Figure 1:**
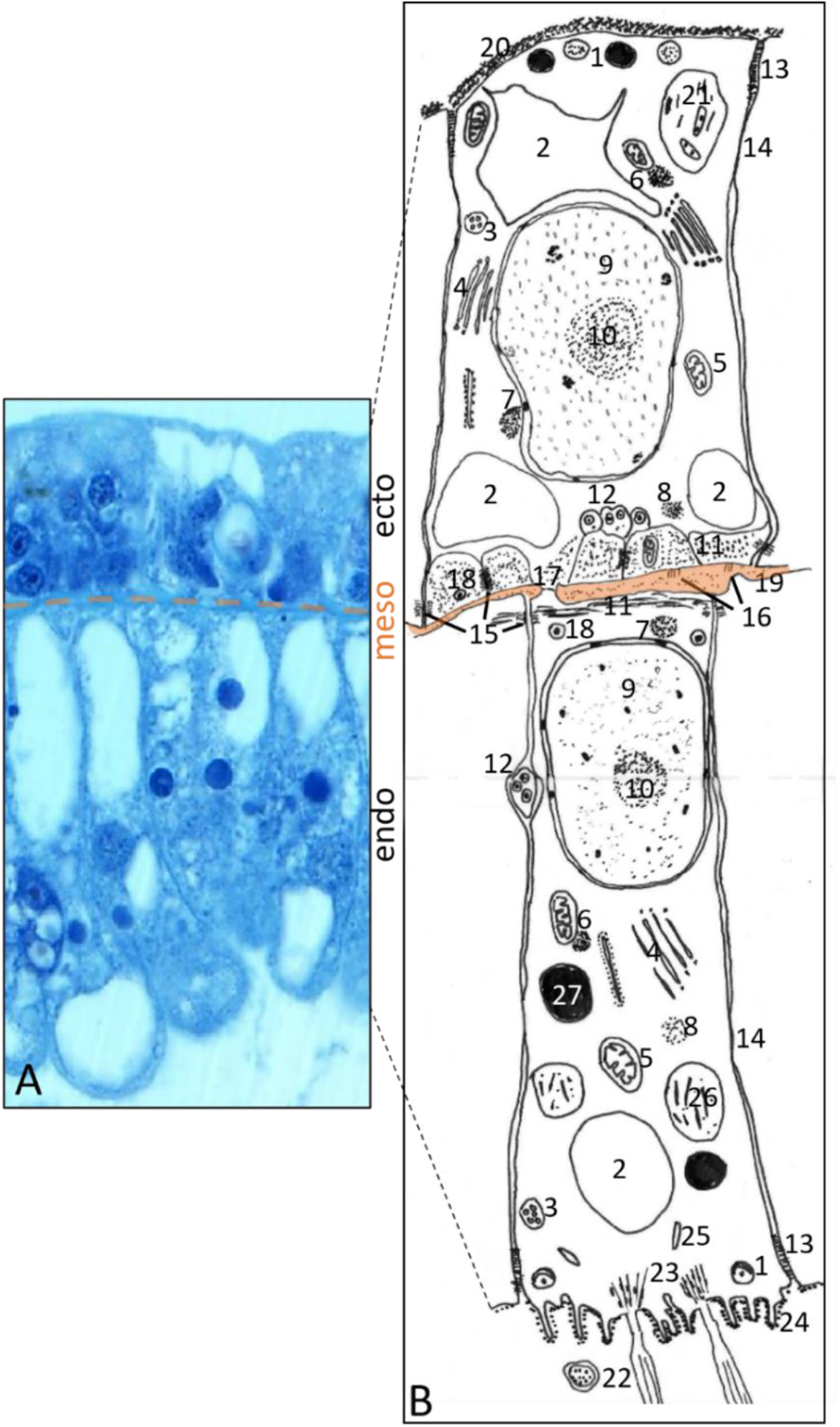
A: LM of cross-sectioned ecto- and endodermal epithelial cells. B: Structures in both layers: 1) apical secretory granules, 2) intracellular pleiomorphic channel system, 3) multivesicular endosome, 4) Golgi stack, 5) mitochondria, 6) nuage (attached to mitochondrium), 7) nuage (attached to nuclear pore), 8) nuage (“free-swimming” in cytoplasm), 9) nucleus, 10) nucleolus, 11) myoprocesses, 12) neurites with dense core vesicles, 13) septate junction, 14) gap junction, 15) desmosome like junctions, 16) hemidesmosome like junctions, 17) gap junction connecting ecto- and endoepithelial cell processes, 18) epithelial dense core vesicle, 19) mesoglea. Ectodermal cell specific structures: 20) cuticle, 21) phagosome with bacteria and cuticle remnants. Endodermal cell specific structures: 22) flagellae, 23) rootlets of flagellae, 24) microvilli with faint glycocalyx, 25) rod shaped bodies, 26) digestive vacuole, 27) lipid droplet.

### Stemness: Epithelial cells exhibit nuclei typical for proliferating cells and nuage

Ultrastructural analysis of ecto- and endodermal epithelial cells revealed conserved nuclear features associated with stemness (Fig. 2). Both epithelial layers exhibited nuclei enriched in non-condensed chromatin (Fig. 2B,H) and prominent nucleoli (Fig. 2C,I). Numerous distinct nuage types - electron-dense, perinuclear structures associated with RNA regulation (Juliano et al. 2014) - were observed, positioned freely in the cytoplasm (“free swimming”) or attached to nuclear pores or mitochondria (Fig. 2D-F, J-L). Mitotic cells remained integrated within the epithelial layer, retaining intact apical septate junctions during mitosis, but disassembled basal muscle processes and the contact to the mesoglea (Fig. 2M). Homologous chromosome pairs and surrounding mitochondria were visible during cell division, highlighting the maintained epithelial organization throughout mitosis.

**Figure 2:**
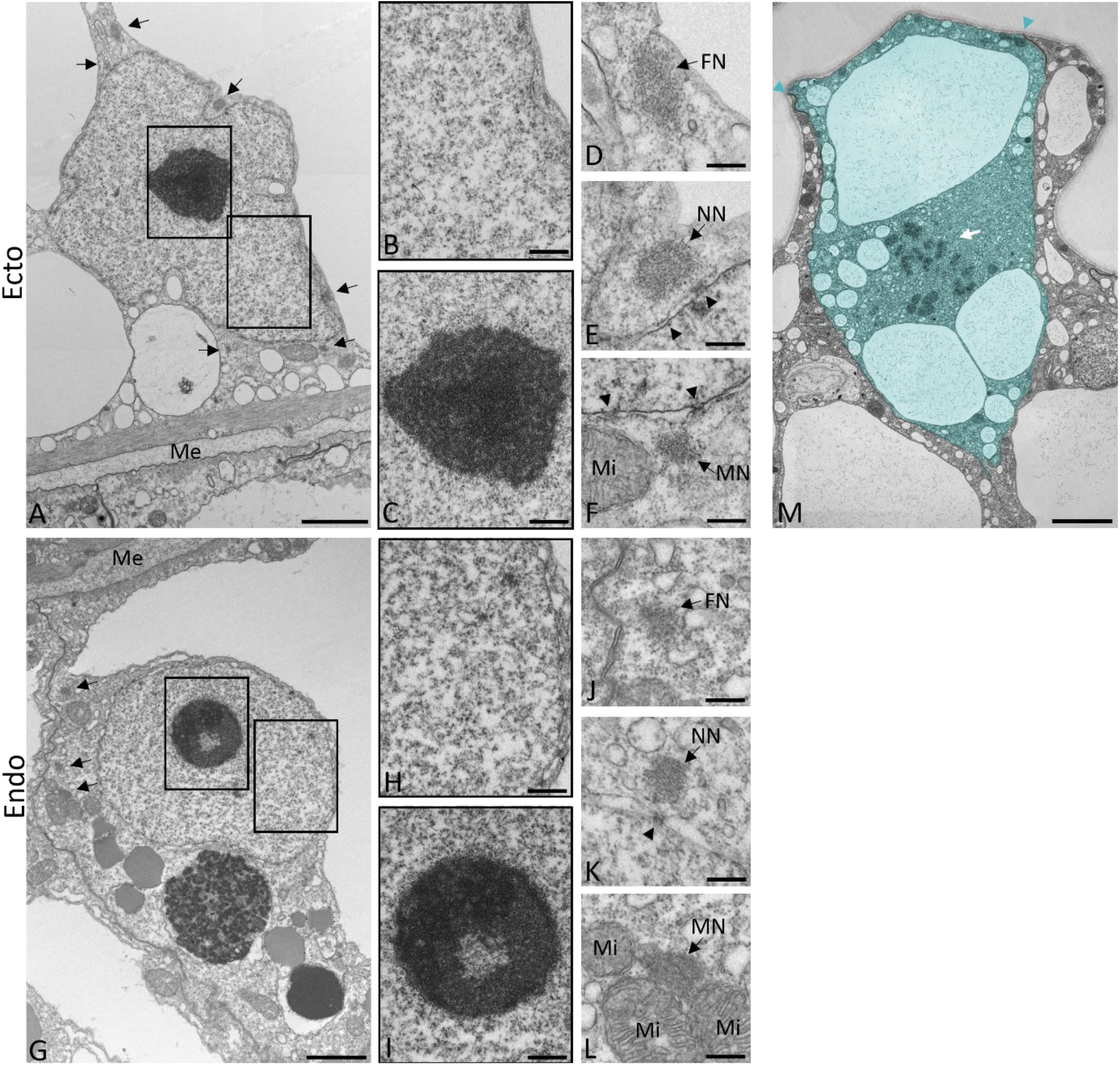
Ecto- and endodermal epithelial cell with nucleus, nucleolus, nuages (A,G) and mitosis (M). Ectodermal (B-F) and endodermal (H-L) cell details of non-condensed chromatin (B,H), nucleolus (C,I), different nuages (D-F,J-L; black arrows; FN = “free swimming”, MN = mitochondrion attached, NN = nuclear pore attached) and nuclear pores (black arrow heads). M) Ectodermal cell in metaphase with intact septate junctions (cyan arrow heads), homolog chromosome pairs (white arrow) and lack of basal muscle processes. Me = Mesoglea, Mi = mitochondria. Scale bars: A,G,M, 2μm; B,C,H,I, 500nm, D-F,J-L, 250nm.

### Differentiated state: Epithelial cells show extensive ultrastructural specializations for transport, secretion, and communication

#### Cryofixation reveals new insights into the pleiomorphic channel systems

In contrast to stem cells, differentiated epithelial cells show extensive channel systems in both ecto- and endodermal layers (Fig. 3A,B). Cryofixation revealed an intracellular, pleiomorphic channel architecture not visible with chemical fixation, suggesting that conventional methods often collapse these structures into large vacuoles (Supp. Fig. 1). These channels may temporarily open to the apical cell membrane being possibly involved in transport across other cell compartments as well as inter- or extracellular space (Fig. 2C). Along, and frequently in contact with the apical cell membrane there also occurred numerous, ∼400nm wide vesicles containing scarce, faintly stained fibrous material, but their nature and possible relationships to other endomembranes are unclear. Macropinocytosis - an endocytic strategy involved in uptake of fluids and nutrients and membrane remodeling – was observed at the apical cell membrane, presenting as actin-supported, cup-shaped cell protrusions (Fig. 3E). Furthermore, live imaging using LifeAct-GFP *Hydra* showed pulsatile actin rings at the apical cell membrane, reflecting macropinocytotic activity (Fig. 3D, Skokan et al. 2024).

**Figure 3:**
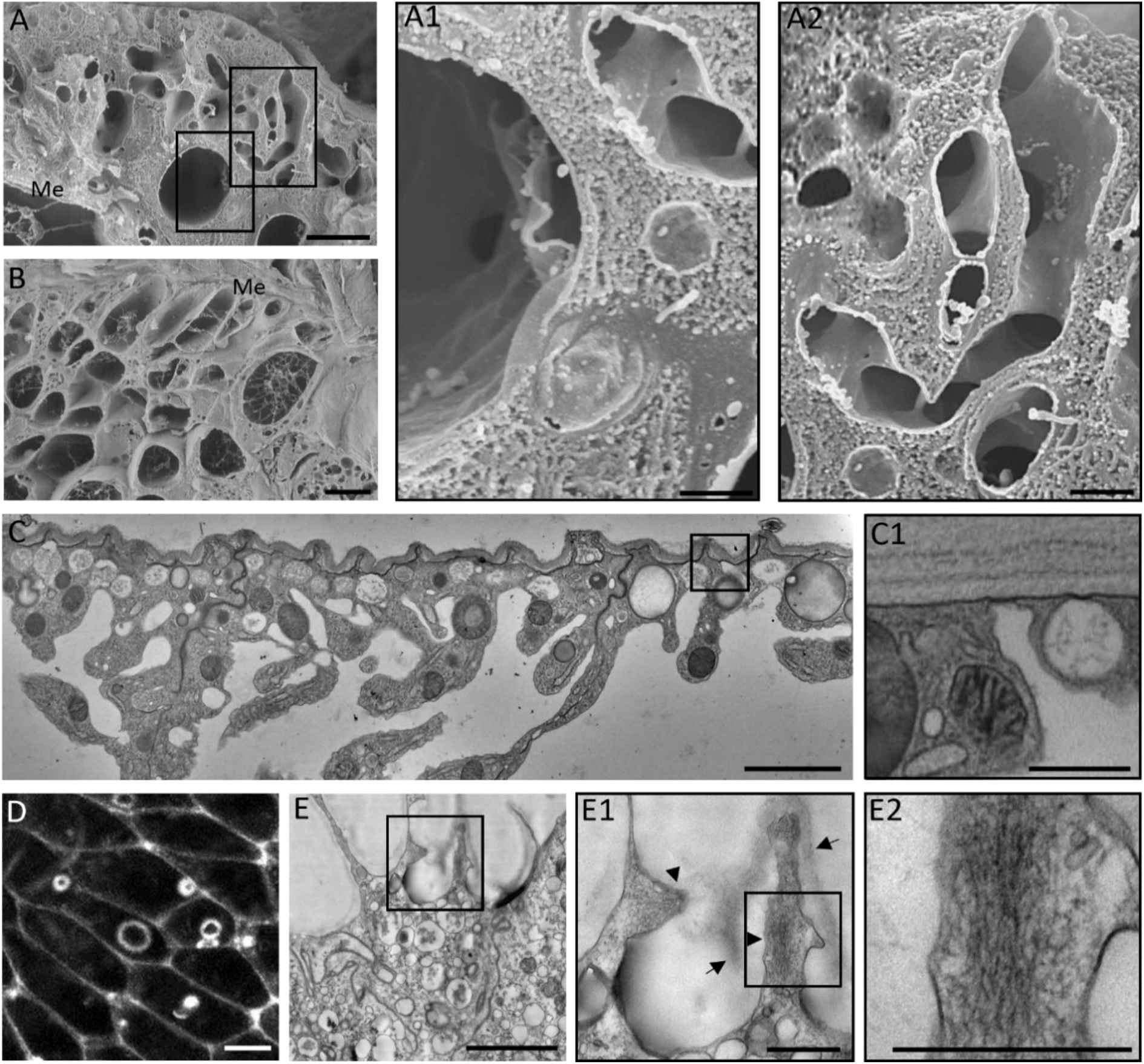
Channel system and macropinocytosis. A-C: Ecto- and endodermal channel system (A,C, and B, respectively) in high-pressure frozen and freeze-substituted *Hydra* as shown by SEM (A,B) and thin section TEM (C). D,E: Macropinocytosis. D) Macropinocytic cups in the ectoderm of a live, LifeAct-GFP-expressing *Hydra*. E) Tangential section through the apical surface of an epithelial cell. E2-3) Detail showing a macropinosome cup with its mesh-like actin core (arrowheads) and the cuticle (arrows). Scale bars: A,B,C,E, 2μm; D, 10μm, A1-2,C1,E1-2, 500nm.

#### Epithelial polarity

The differentiated state is further reflected by the epithelial configuration - the apical-basal and planar epithelial polarity. In *Hydra* the apical-basal polarity is defined by apical specializations such as the ectodermal cuticle, endodermal flagellae/microvilli and apical septate junctions (SJ), as well as by basal-lateral specializations such as basal desmosome-like junctions (DLJ) and hemidesmosome-like junctions (HDLJ) and basal muscle processes. The planar polarity is defined by the orientation of basal muscle processes, with longitudinal ecto- and perpendicular endodermal muscle processes. The *Hydra* junctional complex consisting of ecto- and endodermal apical SJ, gap junctions, basal DLJ and ectodermal HDLJ is shown in Supp Fig. 2 and has been previously described (Chapman et al. 2010; Seybold et al. 2016).

#### Apical specializations for secretion and extracellular matrix production

The apical surfaces of *Hydra* ecto- and endodermal epithelial cells display distinct, specialized ultrastructural features (Fig. 4). The ectodermal apical cell membrane is covered by an up to 1.5μm thick fibrous cuticle formed by 5 distinct layers containing 3 main components - glycosaminoglycans, SWT and PPOD proteins (possibly acquired by horizontal gene transfer from bacteria (Böttger et al., 2012). The proteins are stored in apical secretory granules underneath the cuticle and secreted by the ectodermal cells. The endodermal apical cell membrane bears, in contrast, a faint fibrous glycocalix also surrounding microvilli and flagellae that are anchored by rootlets. In endodermal epithelial cells we also observed partially darkly stained apical vesicles, possibly also representing some kind of secretory granules.

**Figure 4:**
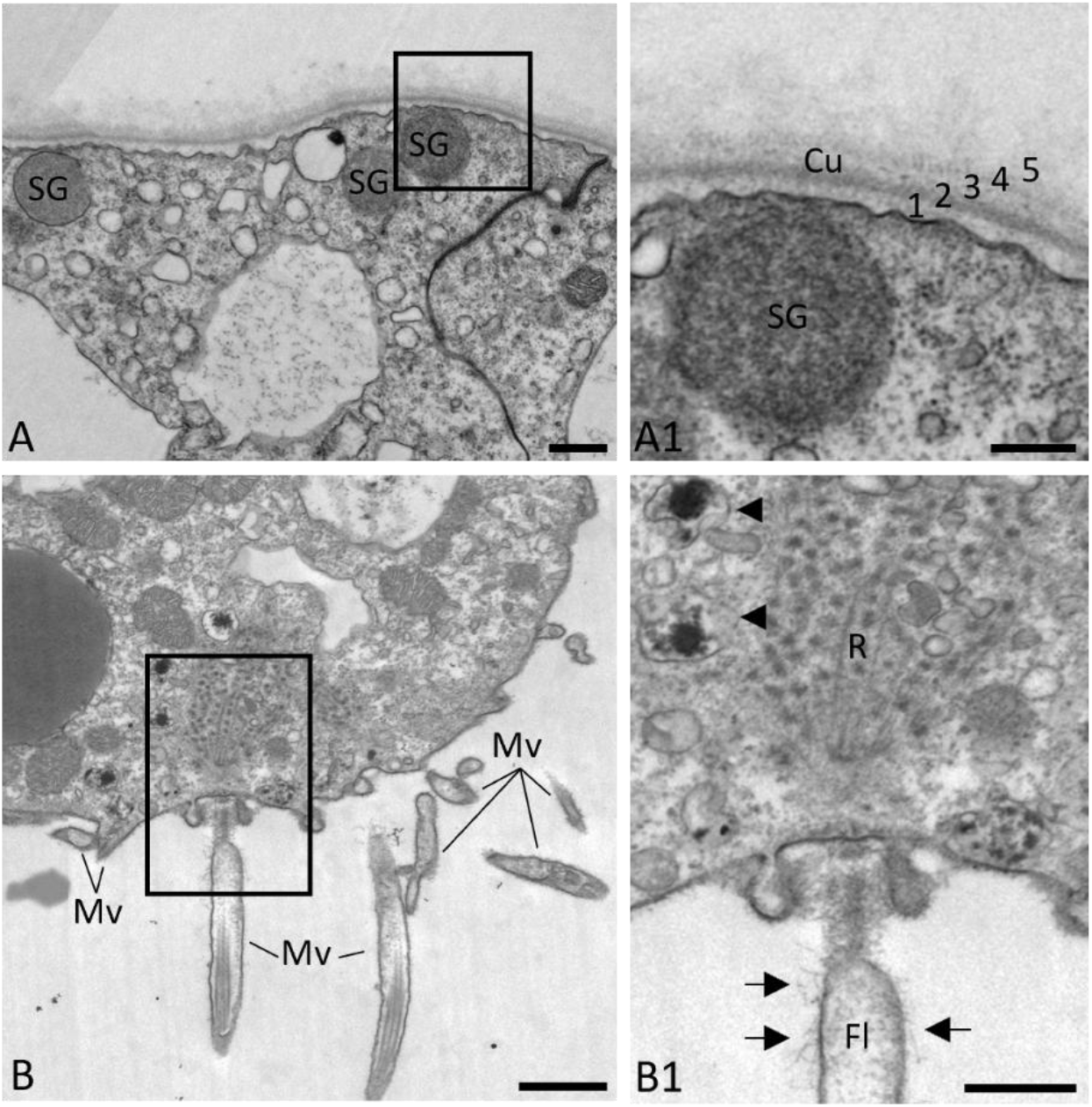
Apical surface of ecto- and endodermal epithelial cells (A and B, respectively. A1) Detail of ectodermal apical surface with cuticle layers (Cu) and apical secretory granules (SG). B1) Detail of endodermal apical surface with flagellum (FI), rootlet of flagellum (R), microvilli (Mv) with faint glycocalix, and apical granules with dark contents (arrowheads). Scale bars: A,B, 1μm; A1,B1, 0.5μm.

#### Intracellular trafficking and vesicle processing

Intracellular trafficking refers to the transport of vesicles and organelles within cells, which is crucial for maintaining cellular functions such as metabolic homeostasis, protein synthesis, recycling, and membrane maintenance in differentiated cells. In *Hydra* endocytic pathways have been analyzed using gold nanoparticles (Marchesano et al. 2013). Nanoparticles (and likely bacteria, cuticle fragments) are rapidly internalized via macropinosomes/phagosomes, recruited and subsequently sorted into different kinds of endosomes to locate finally inside lysosomes (digestive vacuoles/storage vacuoles). This intracellular trafficking is a strong marker for differentiated cells, as it is apparently absent from stem cells (though micropinocytosis may occur in the latter as well).

Using conventional electron microscopy most compartments of intracellular trafficking can be seen, such as ectodermal phago(lyso)somes typically containing bacteria and cuticle remnants, multivesicular endosomes (MVE) and lysosomes/endodermal digestive vacuoles (DV) with (partially) degraded contents (Fig. 5A-E). In the endoderm we further observed “rod-shaped bodies” (Lentz and Barrnett 1965), in addition to above mentioned darkly stained apical granules (Fig. 5D,F).

**Figure 5:**
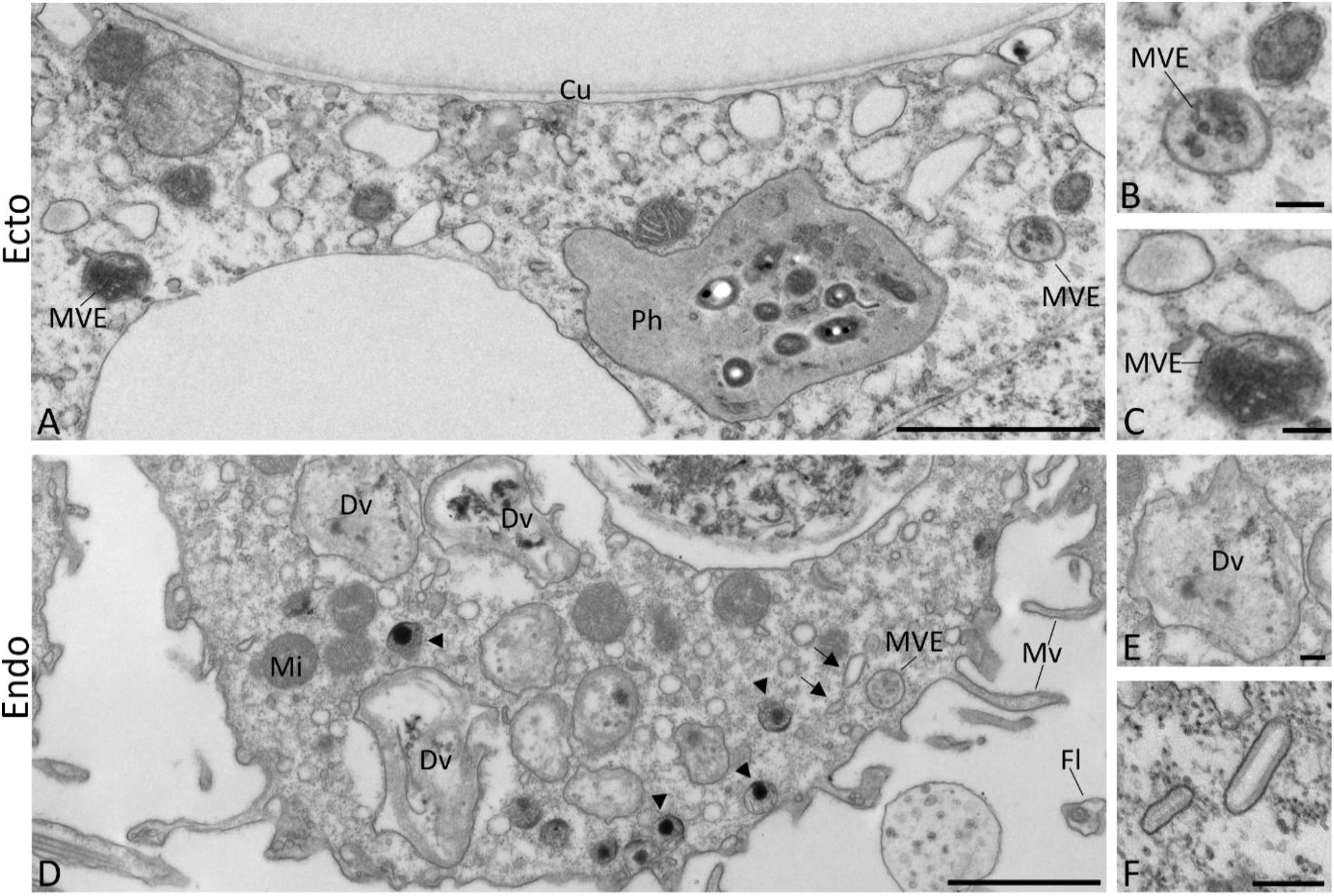
Intracellular trafficking. A) Ectodermal apical surface showing a phagosome (Ph) with bacteria and cuticle remnants, multivesicular endosomes (MVE). B,C) Details of MVEs. D) Endodermal apical surface showing digestive vacuoles (Dv) with degrading contents (E), as well as a multivesicular endosome. “Rod-shaped bodies” (arrows, F) with internal membrane coating are also displayed. Cu = cuticle, Fl = Flagellum, Mi = mitochondrion, arrowheads = dark apical granules. Scale bars: A,D, 2μm; B,C,E,F, 250nm.

Those data on intracellular endomembranes and trafficking are confirmed and refined through use of cryofixation (Fig. 6). In particular, we documented, for example, immature secretory granules in the ectoderm (Fig. 6A,B), in addition to the highly pleomorphic channel system and other vesicles. In the endodermal epithelium the extension and the complex, anastomosing configuration of the channel system became evident (Fig. 6C). Apical granules that frequently contacted the plasma membrane appeared completely filled with darkly stained contents. Finally, we observed a variety of electron-lucent subapical vesicles and tubules (Fig. 6D). Some of them are endocytic (Lentz and Barrnett 1965). Others, including “rod shaped bodies” presumably perform recycling and/or secretion tasks as do similar-looking structures in various mammalian epithelia (Sawaguchi et al. 2004; McDermott et al. 2002; Hudoklin et al. 2011).

**Figure 6:**
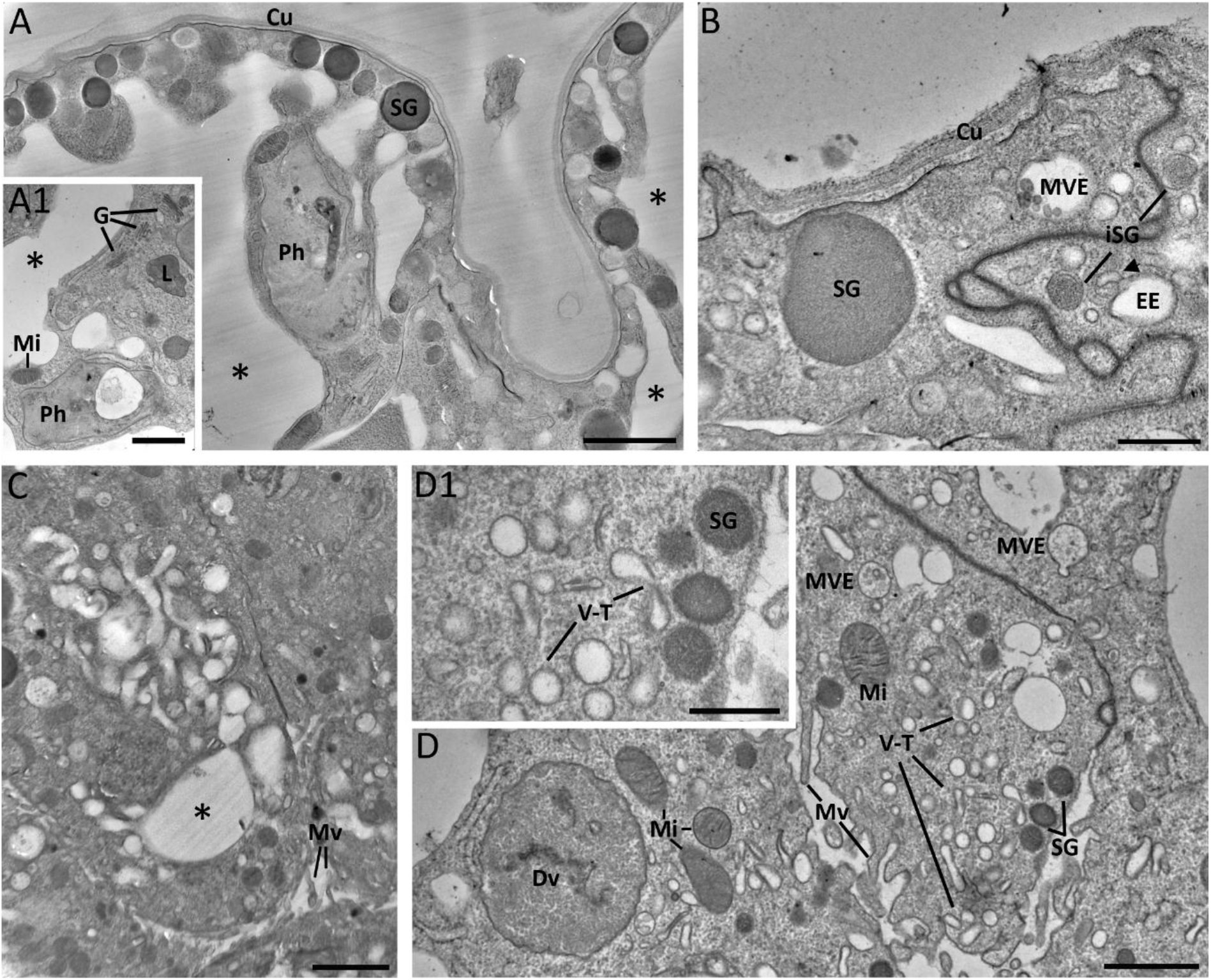
Apical specialization (A-B) and intracellular trafficking (C-D) after cryofixation and freeze substitution of *Hydra*. Cu = cuticle, G = Golgi stack, Dv = digestive vacuole, arrowhead = (putative) recycling tubule of early/sorting endosome (EE), iSG = immature SG, L = lysosome, Mi = mitochondria, Mv = microvilli, MVE = multivesicular endosome, Ph = phagosome, SG = secretory granules, V-T = electron-lucent vesicles/tubules (including „rod shaped bodies”), asterisks = channels. Scale bars: A,C, 2μm; A1,D, 1μm; B,D1, 500nm.

#### Basal specializations and hormone activity

Another strong characteristic for the differentiated state of *Hydra* epithelial cells are their basal specializations such as basal muscle processes, which are separated by the acellular mesoglea. Muscle processes in ectodermal cells run longitudinal and in endodermal cells perpendicular to the body axis and define the planar polarity in the freshwater polyp (Fig. 7). Between these basal muscle processes lay neurites with dense core vesicles (Fig. 7B,D). However, we also observed singular dense core vesicles within the cytoplasm of the epithelial muscle processes (Fig. 7A,C), presumably involved in hormonal activity and communication.

**Figure 7:**
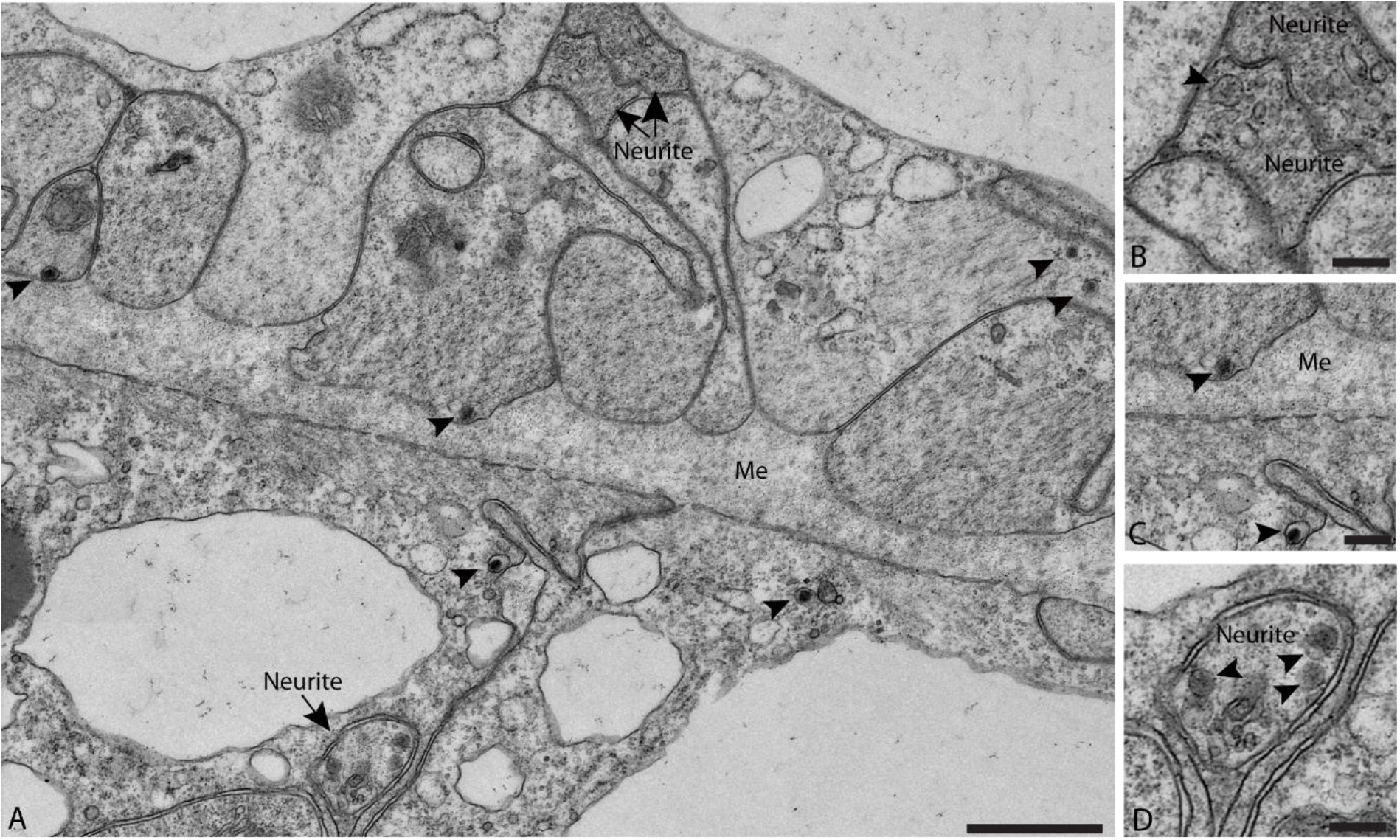
Ecto- and endodermal basal muscle processes with singular dense core vesicles in the cytoplasm (arrowheads), plus associated neurites characterized by numerous dense core vesicles. (B-D) Details showing ecto- and endodermal neurites (B,D, respectively), as well as longitudinal ecto- and perpendicular endodermal actin and myosin filaments, separated by the mesoglea (C). Me = mesoglea. Scale bars: A, 1μm; B-D, 250nm.

## Discussion

Our ultrastructural analysis reveals that *Hydra* epithelial cells occupy a unique position among metazoan epithelia, combining ancient stem cell characteristics with the functional complexity of differentiated tissue. This identity challenges the bilaterian framework in which proliferative and terminally differentiated states are separated into distinct cell lineages. Instead, *Hydra* epithelia maintain proliferative potential while simultaneously performing roles normally assigned to specialized cells, including secretion, transport, contractility, and intercellular communication.

### Stemness vs. differentiation traits in *Hydra* epithelial cells

The stemness features we observe, such as open chromatin, prominent nucleoli, and the presence of nuage, reflect a proliferative program conserved in basal metazoans. Nuages are particularly abundant and enlarged in interstitial cells, correlating with elevated expression of pluripotency markers such as piwi and vasa and their presence in epithelial cells represents the ancestral stem cell machinery (Juliano et al. 2014). Another interesting feature is the epithelial mode of division: cells retain apical junctions during mitosis, maintaining barrier integrity while transiently releasing basal attachments. This behavior shows a proliferative strategy adapted to the constraints of a continuously functioning tissue.

In parallel, differentiated epithelial cells exhibit extensive morphological specialization. The development of apical-basal and planar polarity, the formation of a complex ectodermal cuticle or endodermal glycocalyx, and the presence of microvilli, flagella, and channel systems establish these cells as highly active in secretion, absorption, and membrane trafficking.

Our analysis showed dense core vesicle containing neurites between basal muscle processes, involved in muscle contraction. However, we also observed singular dense core vesicles within the cytoplasm of the epithelial muscle processes, presumably involved in hormonal activity and communication. Takahashi et al. (1997) systematically isolated and characterized *Hydra* peptide signal molecules. These peptides have important regulatory roles e.g. as hormones in endocrine systems, neurotransmitters or neuromodulators in nervous systems or activating factors for lymphocytes in immune systems. In *Hydra* the LWamide peptides (e.g. Hym-54) appear to function as neurotransmitters or neuromodulator, directly inducing muscle contraction. However, these peptides are normally localized in nerve cells and are likely to be absent in epithelial *Hydra* (Takahashi et al. 1997).

### Structure and function of the channel system in *Hydra*

Our ultrastructural analysis including cryo-based EM reveals that *Hydra* epithelial cells possess a far more extensive and complex intracellular channel system than previously known, occurring in both tissue layers. These channels may temporarily open to the apical cell membrane being possibly involved in transport across cell compartments as well as inter- or extracellular space.

Little is known about how water moves and excretion is controlled in ancestral multifunctional epithelia, but is of particular interest due to their evolutionary position between unicellular protozoa with contractile vacuoles and plathelminthes with developed osmoregulatory organs. Osmoregulatory control in freshwater *Hydra* has been investigated by Prusch et al. (1976) Solutes are actively accumulated from the outside across the ectoderm and some of it is secreted into the enteron via the mesoglea and endoderm. Water follows the solute secretion into the enteron, producing a hypoosmotic fluid compared to the body tissue, which gets expelled through the mouth via body contractions, maintaining the tissue of this freshwater organism hyperosmotic to its environment (Prusch et al., 1976).

At the apical cell membrane we observed actin-supported macropinocytotic cups and live imaging using LiveAct-GFP *Hydra* showed pulsatile actin rings (Skokan et al., 2024). Our data indicates, that ectodermal macropinocytosis facilitates uptake of water and solutes from the surroundings, which are then transported through the connecting intracellular channel system and potentially towards the enteron.

### Evolutionary perspective

Tyler (2003) suggested that an embryonic epithelial stem cell is the first distinct cell type to develop during transition from the morula to the blastula stage, and that epithelial stem cells thus likely constitute an ancestral metazoan character. In addition, adult epithelial stem cells seem to be present across extant animal groups (for review see Rinkevich et al., 2022). In sponges, prominent examples of epithelial stem cells are the pinacocytes and choanocytes. Both cell types are building blocks of sponge bodies, integrate multiple physiological functions, contribute to asexual growth and regeneration, and show capacity for somatic and germ line differentiation. Equivalent data are missing for ctenophores, which are regarded together with sponges to represent the earliest branching metazoan phyla. A conclusive proof for an ancestral plasticity of epithelial adult stem cells would require clear evidence for the presence of these cells in ctenophores, and an unambiguous answer, which of the two groups represents the metazoan archetype.

Among cnidarian laboratory models, some hydrozoans including *Hydractinia* show levels of stemness in epithelial cells in the polyp stage comparable to that observed in *Hydra* (Plickert et al., 2012). A notable example of epithelial plasticity is the induction of pluripotency in epithelial cells of *Hydractinia echinata* by experimental activation of the Pou4-related factor Polynem (Millane et al., 2011). For *Nematostella vectensis*, the major anthozoan polyp model organism, evidence that epidermal or gastrodermal epithelial cells act as adult stem cells in dynamic tissue turnover is missing. Nevertheless, we would argue that cnidarian life cycles with features such as extensive, and in some cases indefinite, growth of polyps and polyp colonies, asexual reproduction, and high regenerative capacities would favor the presence of epithelial multi-functionality and plasticity.

To which extent has ancestral epithelial multi-functionality and plasticity been inherited to bilaterians? Actually, only few examples have been described in invertebrates. Among the best-studied examples are epithelial bud primordium cells in colonial tunicates, i.e. in *Botryllus* (see Rinkevich et al., 2022). They serve as a source for a new colony unit during a blastogenic life cycle and differentiate various types of somatic products. High potency is here again linked to asexual colonial growth and longevity. In vertebrates, epithelial multi-functionality is commonly limited to tissue- or organ-specific potency. Several of these epithelial adult stem cells are particularly well-studied, such as stem cells in the intestinal crypts and skin stem cells. Overall, we argue that a division of labor-principle (Arendt et al., 2009) also applies to the evolution of epithelial stem cells, with extensive multi-functionality and plasticity in prebilaterian phyla, and more and more limited multi-functionality and plasticity in bilaterians and vertebrates.

## Conclusion

By examining the ultrastructure of *Hydra* epithelial cells, we highlight a cellular strategy that predates the strict division of labor found in more derived metazoans. This work provides a framework for understanding how epithelial multifunctionality may have shaped early tissue evolution and offers insight into how stemness can be maintained without compromising complex physiological function.

## Acknowledgements

We would like to thank Taylor Skokan for kindly providing the Lifeact images.

## Notes

### Competing Interest Statement

The authors have declared no competing interest.

