## Supplemental Figures for "Ultrastructure of stemness and differentiated state in *Hydra* epithelial cells"

### Supplements

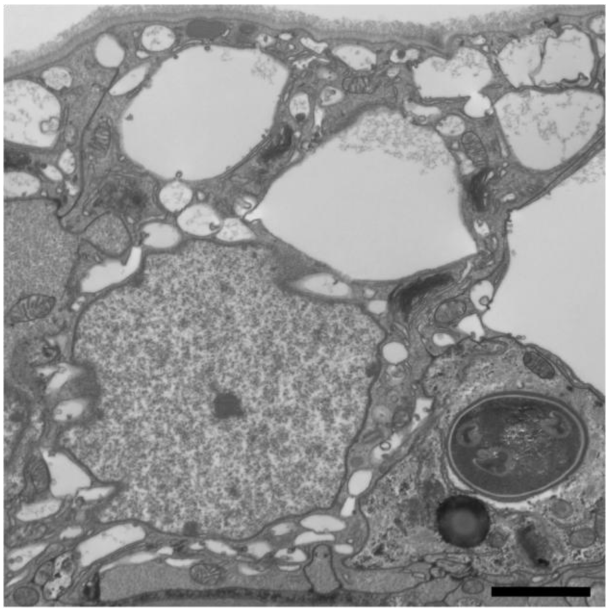

Supp. Figure 1: Ectodermal epithelial cell after chemical fixation, showing large vacuoles instead of a pleomorphic, anastomosing channel system. Scale bar, 2µm.

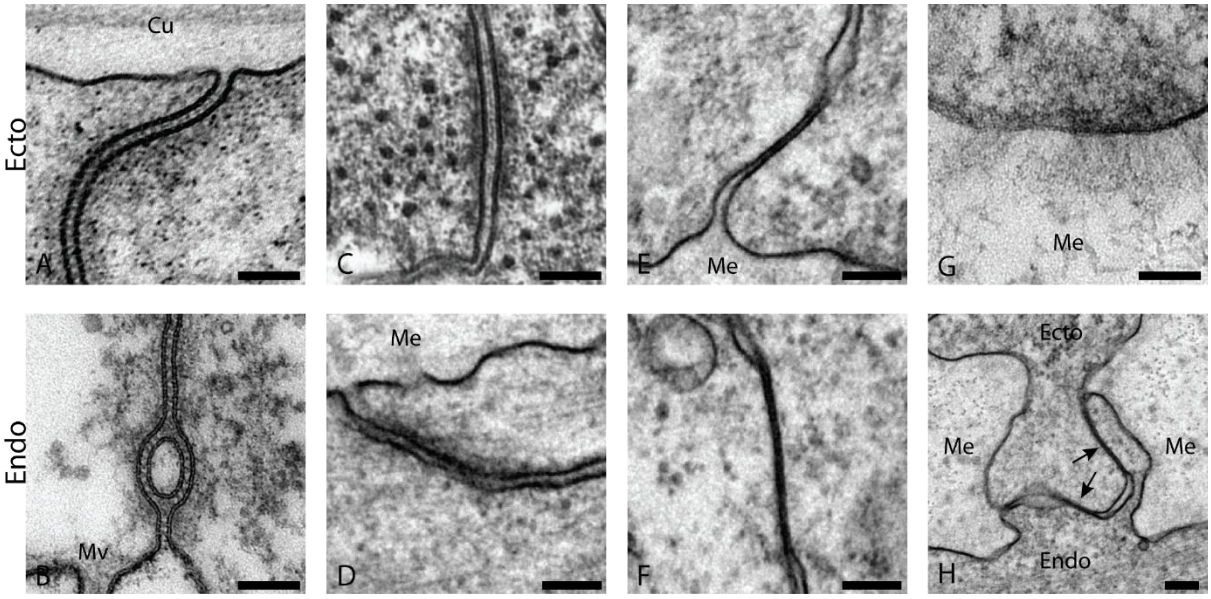


Supp. Figure 2: *Hydra* epithelial junctions: A,B) Ecto- and endodermal apical septate junctions, respectively. C,D) Ecto- and endodermal desmosome-like junctions. E,F) Ecto- and endodermal gap junctions. G) Hemidesmosome-like junction. H) Gap junction between ecto- and endoderm across the mesoglea. Cu = cuticle, Me = mesoglea, Mv = microvilli. Scale bars, 100nm.

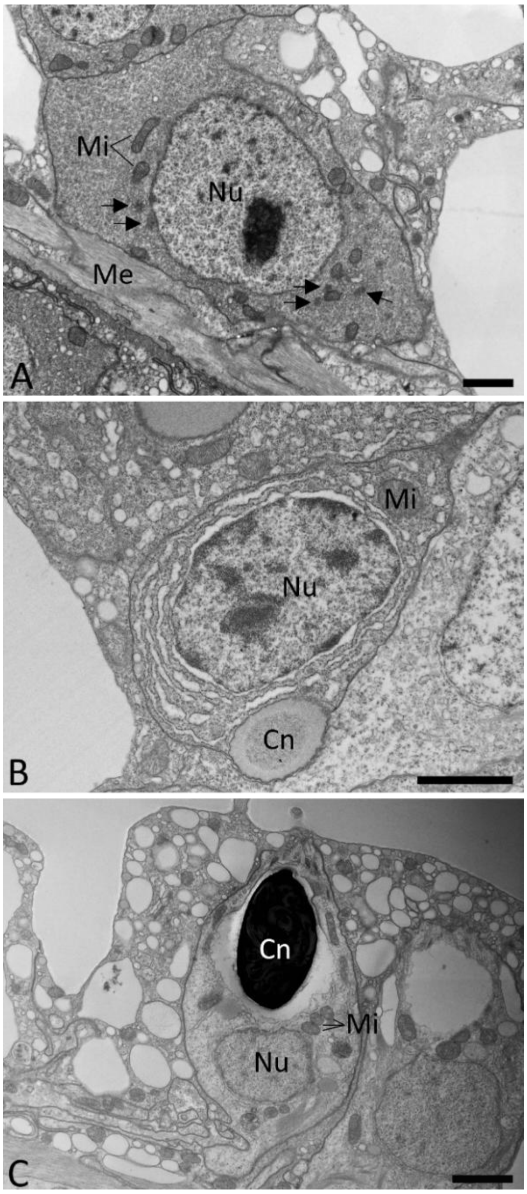


Supp. Figure 3: *Hydra* interstitial stem cell (A) with nucleolus surrounded by non-condensed chromatin and nuages (arrows); differentiating nematocyte (B) with mixed eu- and heterochromatin and differentiated cytoplasm; and fully differentiated nematocyte (C). Cn = Cnidocyte, Me = mesoglea, Mi = mitochondria, Nu = Nucleus. Scale bars, 2µm.
